# The diversity of interaction types drives the functioning of ecological communities

**DOI:** 10.1101/411249

**Authors:** Vincent Miele, Christian Guill, Rodrigo Ramos-Jiliberto, Sonia Kéfi

## Abstract

Ecological communities are undeniably diverse, both in terms of the species that compose them as well as the type of interactions that link species to each other. Despite this long-recognition of the coexistence of multiple interaction types in nature, little is known about the consequences of this diversity for community functioning. In the ongoing context of global change and increasing species extinction rates, it seems crucial to improve our understanding of the drivers of the relationship between species diversity and ecosystem functioning.

Here, using a multispecies dynamical model of ecological communities including various interaction types (e.g. competition for space, predator interference, recruitment facilitation), we studied the role of the presence and the intensity of these interactions for species diversity, community functioning (biomass and production) and the relationship between diversity and functioning.

Taken jointly, the diverse interactions have significant effects on species diversity, whose amplitude and sign depend on the type of interactions involved and their relative abundance. They however consistently increase the slope of the relationship between diversity and functioning, suggesting that species losses might have stronger effects on community functioning than expected when ignoring the diversity of interaction types and focusing on feeding interactions only.

## 1 Introduction

Despite the wide recognition of the coexistence of multiple interaction types linking species in nature [1, 2, 3], research on ecological networks has been massively dominated by studies on a single interaction at a time (e.g. trophic, competitive or mutualistic; e.g. [4, 5, 6]). The implications of the diversity of interactions for ecological community dynamics and resilience remains therefore largely unknown, despite a recent growing interest in the ecological literature [7, 8, 9, 10].

Among interaction types, feeding has massively dominated the literature [2], leading to the analysis of the structural properties of food webs on data sets and to the use of modeling to investigate the functional consequences of these structures (e.g. [4, 11, 12, 13, 14, 15, 16]). Early on, Arditi and colleagues [17] proposed to integrate non-trophic interactions in such dynamical models as modifications of trophic interactions (so-called ‘rheagogies’). Building on that idea, Goudard and Loreau [18] investigated the effect of rheagogies on the relationship between biodiversity and ecosystem functioning (BEF) in a tri-trophic model. They showed that ecosystem biomass and production depended not only on species richness but also on the connectance and magnitude of the non-trophic interactions.

Several studies have investigated the role of incorporating specific interactions in food webs. For example, incorporating interspecific facilitation in a resource-consumer model allowed species coexistence in communities of plants consuming a single resource [19]. This increase in species diversity also happens in ecological communities with higher trophic levels including both trophic and facilitative interactions [3]. In the same model, intra- and inter-specific predator interference increased species coexistence as well in multi-trophic webs, although to a lesser extent than facilitation among plants [3].

More generally, the joint effect of several interaction types is expected to affect community functioning and stability. Extending May’s work, Allesina and Tang [20] showed that communities including the same amount of positive and negative interactions were less likely to be stable than random ones (i.e. where interactions between species are randomly chosen), themselves being less stable than predator–prey communities. In a spatially explicit model including both mutualism and antagonism, Lurgi *et al.* [9] found that increasing the proportion of mutualism increased the stability of the communities. Addressing the relationship between structure and stability, Sauve *et al.* [8] showed that the role of nestedness and modularity – structural properties that are known to promote stability in single interaction types networks – was weakened in networks combining mutualistic and antagonistic interactions. Combining dynamical models with an empirical network analysis including all known non-trophic interactions between the species of intertidal communities in central Chile [21], Kéfi *et al.* [10] found that the specific ways in which the different layers of interactions are structured in the data increased community biomass, species persistence and tend to improve community resilience to species extinction compared to randomized counter-parts. More recently, García-Callejas *et al.* [22] used a dynamical model to investigate the effect of the relative frequency of different interaction types on species persistence and showed that persistence was more likely in species poor communities if positive interactions were present, while this role of positive interactions was less important in species-rich communities.

Altogether, these studies suggest that the joint effect of several interaction types could alter fundamental properties of ecological systems – such as species coexistence, production and community stability – with however a clear lack of consensus on how. So far, most studies have addressed these questions with specific subsets of non-trophic interactions [3, 8, 18, 19], in small species modules [23, 24], in networks with limited numbers of trophic levels [19] or with unrealistic trophic structure [18]. Only a few studies have extended these approaches to complex networks of interactions with a diversity of interaction types (see e.g. [9, 22, 25]). We therefore still lack a clear view on the overall role of the diversity of interaction types *per se* for species diversity and community functioning, and especially how they may affect the relationship between diversity and functioning.

In the 90ies, because of the raising awareness of the increase in species extinction rates, the longlasting interest on the origin and maintenance of species diversity shifted toward the study of the consequences of biodiversity, and especially of its loss, for ecosystem functioning [26]. This became an entire sub-field of ecology referred to as ‘Biodiversity and Ecosystem Functioning’ (so-called BEF) and lead to decades of experimental and theoretical research investigating how diversity affects functioning (see [27, 28, 29, 30, 31, 32] for reviews). Results of experimental studies suggests that more diverse communities generally produce more biomass than less diverse ones [33, 34]. Theoretically, the question has been addressed as well; models have long focused on plant communities (i.e. a single trophic level) (e.g. [35]), but have more recently started to expand these investigations to more complex, realistic communities (e.g. [36, 37, 38]). Until now, as far as we know, studies had not specifically investigated the role of the diversity of interactions types on the shape of the BEF.

Here, using a bioenergetics resource-consumer model in which broad categories of non-trophic interactions were introduced [3], we systematically investigated the functioning of ‘multiplex’ ecological networks, i.e. how multiple interactions (their abundance and intensity) affect species coexistence, community functioning (biomass and production), and the relationship between diversity and functioning. Our model includes, in addition to the consumer-resource interactions, competition for space among sessile species, predator interference, refuge provisioning, recruitment facilitation as well as effects that increase or decrease mortality.

## 2 Results

In what follows, we use NTI(s) to refer to non-trophic interaction(s).

### 2.1 Effect of the presence and intensity of each NTI

We ran community dynamics with or without NTIs, and evaluated the relative difference in community characteristics at steady state obtained in the presence compared to in the absence of each NTI. This allowed comparing the effects of the different NTI types (for a range of interaction intensities; see Methods).

We found that interference had a negative effect on diversity and community production and a weak (positive) effect on biomass (1st row of Fig. 1). Through time, interference decreased the consumption of some predators (those having prey in common); this initially favored some of the basal and intermediate species (that were less consumed), and eventually lead to the extinction of some of the intermediate and top predators. The remaining consumers, relieved from competition for their prey, gained biomass, which compensated for the losses due to the extinctions (see supplementary Fig. A.1 and A.2).

**Figure 1:**
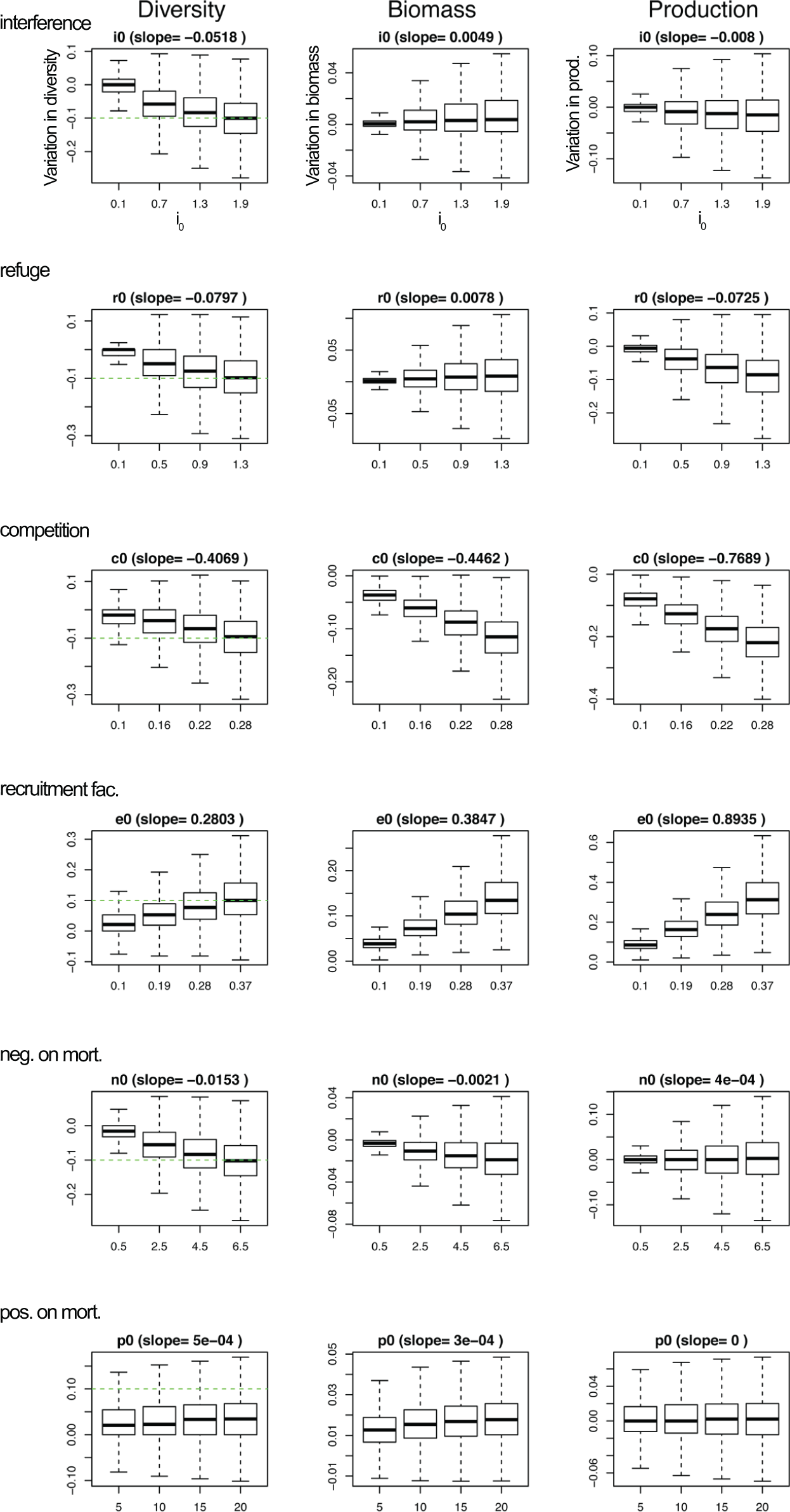
Variations in species diversity, biomass and production (in columns) as a function of the intensity of the NTI along the x-axis for each of the 6 NTIs (in rows). Values along the y-axis are evaluated at steady state in networks with NTIs compared to ones without NTIs. Values on the x-axis correspond to values of the parameters *i*_0_ for interference, *r*_0_ for refuge provisioning, *c*_0_ for competition for space, *e*_0_ for recruitment facilitation, *n*_0_ for increase in mortality (i.e. negative effects on mortality), *p*_0_ for decrease in mortality (i.e. positive effects on mortality). For each NTI, their maximum value along the x-axis was chosen such that it lead to a ±10 % change in species diversity relatively to the case without NTI (green dashed line). Note that the y-axis of the different panels differ. The NTIs were categorized into ‘positive’ (i.e. beneficial, recruitment facilitation) vs ‘negative’ (i.e. detrimental; interference, refuge provisioning, competition for space and increase in mortality) based on their effect on diversity.

Refuge provisioning had similar overall effects, but its negative effect on production was about 9 times larger than the one of interference (2nd row of Fig. 1). In this case, species benefiting from refuges remained in the system but were less accessible resources. This lead to a loss of biomass and subsequent extinctions of some top predators (which could not access their prey), but the ones remaining in the system did not benefit from these extinctions, since most of the resources were still under protection (except for those whose protector went extinct). The loss of diversity was in this case not entirely compensated by a gain in biomass as for interference (see supplementary Fig. A.1 and A.2, 2nd row).

Competition for space had a negative effect on all variables, while facilitation for recruitment had a positive effect on all community characteristics (3rd and 4th rows of Fig. 1). Through time, these effects first affected the basal species, then the intermediate and eventually the top predators (see supplementary Figs. A.1 and A.2).

Modifications of mortality rates produced very weak effects overall. Increasing mortality had a negative effect on diversity and biomass and a very weak effect on production (3rd row of Fig. 1). Decreasing mortality had a weak positive effect overall. We did not consider these last interactions in what follows and focused instead on the five remaining NTIs, namely interference among predators, refuge provisioning, recruitment facilitation, competition for space and increase in mortality.

Overall, the most influential NTIs among the ones studied were competition for space and facilitation for recruitment, in terms of both community diversity and functioning (see slopes linking the parameter values to see the extent of the effects in Fig. 1). The effects of competition for space on diversity were about 8 and 5 times larger than those of interference and refuge, while the effects on biomass were resp. 91 and 57 times larger. Regarding recruitment facilitation, the effects on diversity were about 5 and 3 times larger than those of interference and refuge, and the effects on biomass were resp. 78 and 49 times larger.

For all NTIs, effects seemed to be stronger on intermediate and top trophic levels at steady state (see supplementary Fig. A.2). Regarding species diversity, this was partly due to the fact that plant species already all persisted with trophic interactions only, therefore NTIs had more leverage on intermediate and higher trophic levels where species did not all persist in webs with trophic interactions only. Regarding biomass, effects seemed to first affect basal species but then climb up the food web to eventually affect mainly the top predators.

This first set of simulations helped us categorize the NTIs studied into ‘positive’ (i.e. beneficial; recruitment facilitation) vs ‘negative’ (i.e. detrimental; inter-specific predator interference, refuge provisioning, competition for space and increase in mortality) based on their effect on diversity.

### 2.2 Combined effects of the NTIs on species diversity

We now mixed the five remaining NTIs together, with NTI intensities picked at random within predefined ranges (see Methods and Fig. 1) to study the joint effect of the NTIs considered.

Not unexpectedly, the effect of the presence of the NTIs depended on the relative number of links of the different NTIs and on their intensities. When all interaction types were together with an equal proportion of positive and negative NTIs, networks with NTIs tended to have a smaller species diversity than networks without NTIs (Fig. 2A). In other words, NTIs lead to extinctions of species compared to simulations run with feeding interactions alone. There were also quite a few number of cases where the net effect on diversity was null.

**Figure 2:**
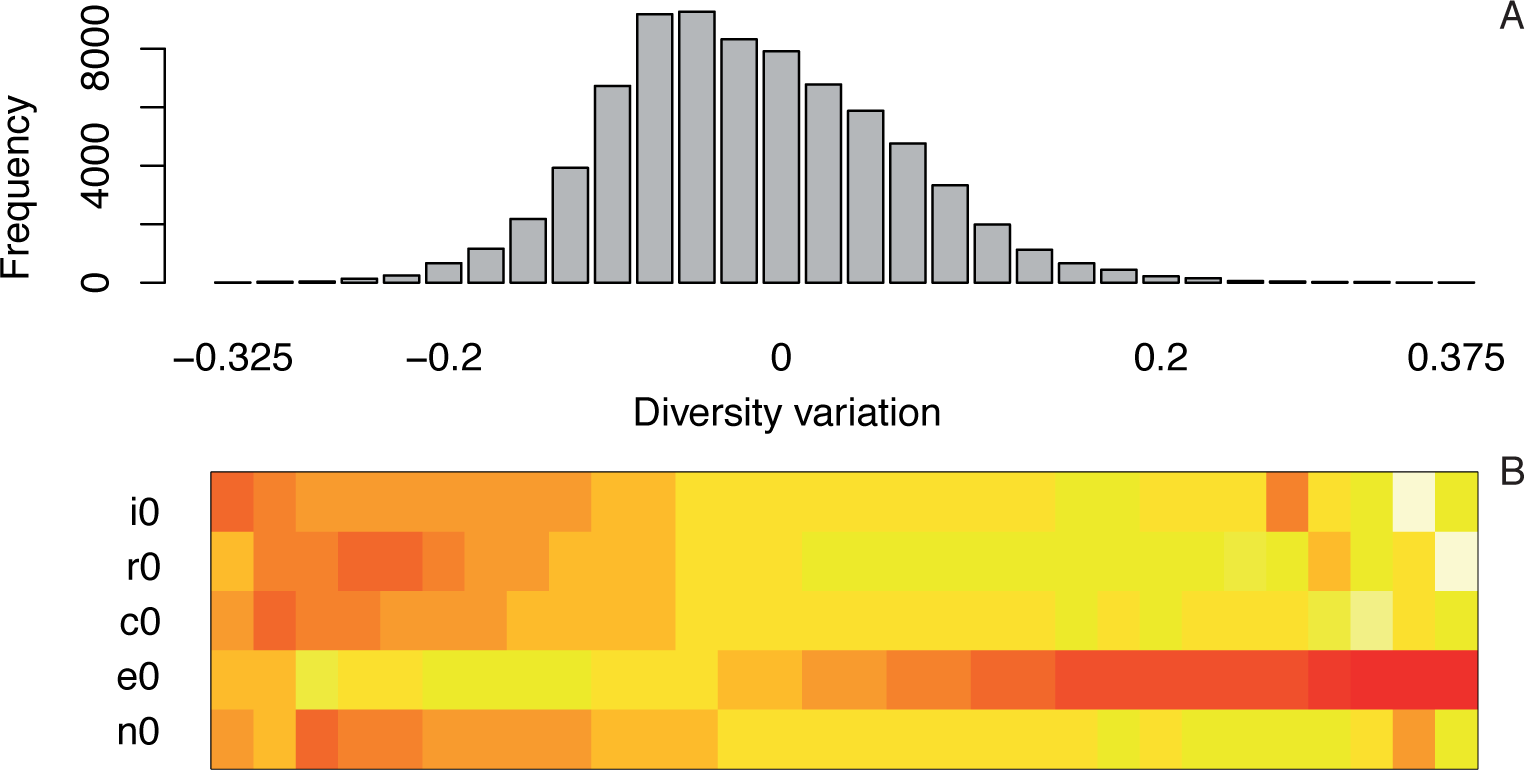
A) Frequency of simulations leading to a given variation of diversity (x-axis) in networks with trophic links only compared to those with both trophic and non-trophic links. In these simulations, NTI intensities were picked randomly at the start of the simulation in the ranges defined in Fig. 1. B) Average values of each of the NTI parameters corresponding to the simulations of A (mean parameter value used for the simulations leading to each of the bar in A). Colors range from light yellow for small average values to red for strong average values. *i*_0_: intensity of interference among predators, *r*_0_: intensity of refuge provisioning, *c*_0_: intensity of competition for space, *e*_0_: intensity of facilitation for recruitment, *n*_0_: intensity of increase in mortality.

There was nonetheless a fraction of cases where NTIs tended to enhance species diversity; these were clearly cases where beneficial NTIs were present and strong (orange and red areas on Fig. 2B). It was noteworthy that the NTI values were all chosen at random for each of the simulations, so all combinations of intensity values were possible and present across simulations, but our results showed that positive effects of NTIs on diversity always happened when the beneficial NTI (recruitment facilitation) was strong while the detrimental NTIs were weak.

Now fixing the intensities of all NTI links to their maximum value (corresponding to a 10% effect on diversity variation; see Fig. 1) and focusing on their relative abundance, we found that a greater number of recruitment facilitation links tend to favor positive effects on diversity while increasing the number of interference, refuge or competitive links pushed toward negative effects on diversity. (Supplementary Fig. A.3).

### 2.3 Combined effects of NTIs on the Biodiversity-Ecosystem functioning relationship

How did these effects on species diversity translate into community functioning? Using the previous simulations where NTI intensities were picked at random, we found that both in food webs and in ecological networks with NTIs, the relationship between species diversity and biomass at steady state was positive (Fig. 3 A, B; this was also the case for production: see Supplementary Fig. A.4). Strikingly, in presence of NTIs, the relationship was significantly stronger than in their absence (ANCOVA p-value<1e-16; comparing slopes in Fig. 3 A and B).

**Figure 3:**
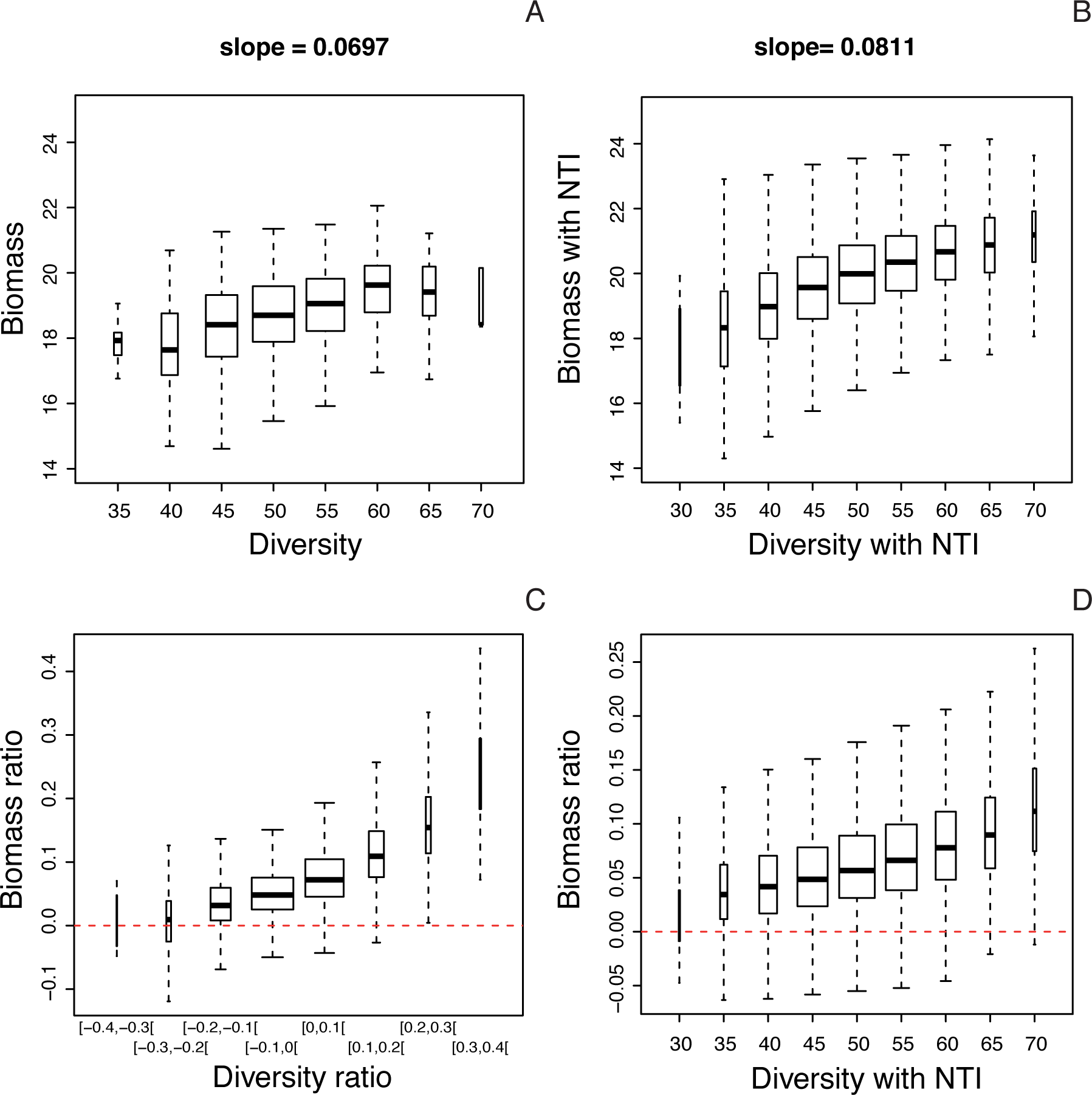
Relationships between species diversity and biomass in networks with compared to without NTIs. A) Biomass as a function of species diversity (number of species) in networks with trophic interactions only. B) Same as A in networks with NTIs (values of all NTIs intensities taken randomly in a given range; see Methods). C) Variation in biomass (in networks with NTIs compared to networks with TI only; see Methods) as a function of the variation in species diversity (in networks with NTIs compared to networks with TI only; see Methods). D) Variation in biomass as a function of species diversity in networks with NTIs.

Plotting the variation in biomass (with NTIs compared to without NTIs) as a function of the variation in species diversity suggested that when NTIs contributed to a gain in species, this generally translated into a gain in biomass as well (Fig. 3 C). Actually, networks with NTIs tended to gain biomass (compared to networks without NTIs) even when there was no gain (or even a weak loss) in diversity (see boxes at diversity ratios of −0.2, −0.1 and 0 in Fig. 3 C). When there was a small loss of diversity in presence of NTI (−0.1 – −0.3), the remaining species took advantage of these extinctions and gained biomass. When there was a gain in species diversity compared to the case without NTIs, this was often happening because of the presence of beneficial NTIs (Fig. 2 B), and those positive links lead to a considerable increase in biomass as well. Note that recruitment facilitation creates by far the strongest diversity-functioning relationships (the strengths of these relationships can be estimated by dividing the slopes for the two measures of ecosystem functioning (biomass and production) by the slope of diversity in Fig. 1). This also explains why NTIs have a positive effect on biomass even when the diversity ratio is zero. These combined effects lead to the positive slope of the relationship between diversity and biomass in the presence of NTIs. There was, however, a large variability around these trends due to the fact that each simulation corresponded to a different combination of NTI intensities.

The gain in biomass in networks with NTIs (compared to the cases without NTIs) seemed to accelerate with species diversity (e.g. species diversity above 50 species in Fig. 3 D).

## 3 Discussion

Using a bioenergetic model in which six types of NTIs were incorporated, we found that these NTIs in isolation and jointly affected significantly species diversity and community functioning (biomass and production), consistently with previous studies addressing the role of the diversity of interaction types in module or network contexts [3, 10, 17, 18, 19, 20, 22].

Overall, when taken together and with a balanced number of beneficial and detrimental interactions (as defined by their individual effects on diversity), the presence of NTIs tended to have a slightly negative effect on species diversity. This is in agreement with Goudard and Loreau [18] who studied NTIs that are modifications of feeding links; with equal numbers of positive and negative effects, they found a decrease in the total number of species when NTIs are incorporated. In our case, this result was not expected since the range of NTI intensities spanned was selected such that each interaction type had equivalent effects on diversity when taken individually. Therefore, despite controlling for both the number and the intensities of NTIs, the joint effect of the NTIs, when simultaneously incorporated in the model, was negative for the species diversity of the resulting communities.

Surprisingly, we found interference between predators to have a negative impact on diversity, which contrasts with other studies reporting stabilizing effects on population dynamics and positive effects on diversity [12, 39, 40] when interference is included via a Beddington-DeAngelis functional response [41, 42]. However, these studies included interference either as a purely intra-specific effect or at least assumed that intra-specific interference was stronger than inter-specific interference. Here, intra-specific interference was explicitly excluded, as it was the aim to study inter-specific NTIs only. In this sense, our result that interference reduces diversity is reflecting classic results from competition theory, namely that competition is destabilizing if it is stronger between species than within species [43]. As interference was only included for predators that are already competing for at least one common prey species, the decline in diversity can be attributed to an increased effect of the competitive exclusion principle [44]. It has to be noted, however, that the complex network structure of trophic and non-trophic interactions provides a plethora of niches, which reduces the direct applicability of this principle [13]. When including intra-specific interference, the results of our model are consistent with these previous studies (Supplementary Fig. A.5).

Interestingly, we found that NTIs affected the relationship between diversity and functioning. Despite the array of possible effects from the NTIs, individually and combined, the relationship between species diversity and biomass was found to be significantly stronger in networks with NTIs than in networks without them. Again, this was not necessarily expected since simulations were run with all NTIs together, whose intensity values were picked at random in a range such that their effects on diversity was controlled; we could therefore have expected, e.g. compensatory or negative effects since most of the NTIs studied here tended to have negative effects on diversity (Fig. 1). The effect of NTIs on the slope of the diversity-biomass relationship means that when species-rich networks gain even more species, that goes with disproportionately more biomass gain in the presence than in the absence of NTI. This also means that, conversely, species-poor communities lose more biomass with additional species loss with than without NTI. This is due to the fact that species-rich communities are communities in which beneficial interactions are present and strong, while species-poor communities are communities where detrimental NTIs operate. This result is interesting in that it suggests that species loss may have stronger consequences on community functioning than expected if ignoring non-feeding interactions.

Of course, our study presents a number of limitations. The strongest NTIs studied here, namely competition for space and facilitation for recruitment, both affect mainly plants in our model. It is therefore unclear whether these NTI types appear to exert stronger effects because plants are the affected species. This could be a topic of further investigations.

Moreover, we have focused on a selection of six NTIs that are the major ones found to occur in the Chilean web [10,21], but other interaction types not present in this data set are known to be frequent and important in nature. Examples are parasitism, effects on resource availability, plant dispersal or animal movement. Further work could introduce these other interactions in a single framework. We also assumed here the intensity of the interactions between pairs of species to be constant through time. Some interactions may however be context-dependent (beyond the biomass or abundance of other species). For example, adaptive inducible defenses are a form of phenotypic plasticity that affects the strength of predator-prey interactions [45]. Changes in morphological, behavioral, or life-historical traits in response to chemical, mechanical or visual signals from predators have been reported in the literature for a number of organisms [46]. These responses can moreover occur with a lag, given that the expression of defenses may involve considerable time, relative to the organisms life-cycle [47]. The intensity, and even the type of interactions, could also change with e.g. changes in abiotic factors such as climate.

We have no information regarding the relative importance or intensities of the different interaction types. We proposed a way of putting all interaction types on equal footing regarding their effect on species diversity. This is however a debatable choice – we could for example have chosen to make NTIs comparable regarding their effect on biomass. In nature, it is likely that interactions intensities are not equivalent and that some of them are much stronger than others. Making progress along these lines requires experimental work aiming at quantifying different interactions types, which involves a number of challenges [1].

In this study, NTIs were plugged in the food web randomly although with a number of constraints based on our knowledge of the Chilean web [10, 21]. We did not explicitly investigate the role of the structure of the NTI network, despite the fact that previous studies have suggested that it might play an important role [10, 48]. This remains complicated since it can be necessary to take into account the dependency between the structure of the different layers (e.g. NTI types, as observed in [10]); however this is a promising avenue of future research.

Previous studies have used the community matrix approach focused on net effects between species to investigate the role of the diversity of interaction types [7, 20]. This approach has a number of advantages, including the fact that it allows analytical predictions and generalizations. Here, we chose to focus on a more mechanistic approach, starting from the mechanism of the NTIs without assuming their net effect. For example, we had initially assumed that refuge provisioning would be a beneficial NTI, meaning that it would have a positive effect on species diversity. We however found the opposite in the model simulations - indeed, refuge provisioning protects prey from their consumers but they also deprive consumers from their resource and it seems that this latter effect has stronger consequences at the community scale. In a dynamical model, Gross [19] showed that interspecific facilitation among plants allowed the maintenance of species diversity despite the fact that the net effects measured among plants remained negative. These results highlight that insights gained from the analysis of few-species systems cannot be easily translated into the dynamics of complex communities. Focusing on the net effects between species may conceal important coexistence mechanisms when species simultaneously engage in both antagonistic and positive interactions and stresses the importance of working with mechanistic models to better understand the consequences of NTIs for community diversity and functioning. Nonetheless, a more mechanistic approach implies, for each of the NTI identified, to model it in one specific way, matching our knowledge of how these interactions operate in nature (in our case here, having the Chilean web in mind [10, 21]). For example, regarding refuge provisioning, alternative ways of how it could affect trophic interactions are certainly conceivable. It could increase the hill coefficient, e.g. to mimic the fact that prey at low density would become better protected, but if prey density is too high, the predators would still see them. This could have a more positive effect on diversity than found here. Moreover, one NTI of this type affects multiple attack rates at once (all predators of the protected prey are affected), which could be one of the reasons why this NTI has such a strong effect.

Our study is a step toward getting a better understanding of the dynamics of multiplex ecological networks (i.e. including several interaction types among a set of species), and more precisely of the role of NTIs on community functioning. Our model results suggest that, when simultaneously included, and assembled according to simple rules reflecting observations in nature, NTIs tend to mechanically strengthen the BEF, making the dependency between the number of species present in the community and the functioning of this community (in terms of biomass or production) stronger. This result has important consequences for predicting the consequences of species loss on community functioning.

## 4 Methods

### 4.1 The dynamical model

#### 4.1.1 The trophic model

We used an allometric-scaling dynamic food web model [12, 49]. These models have been used extensively to explore the dynamics and stability of complex ecological networks [12, 13, 50].

The food web model consisted of plants (primary producers, at the base of the network, which consume nutrients not explicitly modeled here), and consumers (animals which eat plants and other consumers). The number of species (plants and consumers) and the structure of the web (who eats whom) were initially determined based on the niche model [51] (for details see section 4.2). We mapped dynamical equations to that food web skeleton. The change in species *i*’s biomass density *B*_*i*_ (in 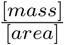)was described by an ordinary differential equation of the general form:

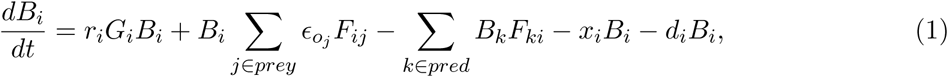

where the first term describes plant growth; the second term describes the biomass gained by the consumption of other species *j*; the third term describes mortality due to predation, summed over all consumers *k* of species *i*; the fourth term represents the metabolic demands of species *i*; the last term is natural mortality of species *i*.

### More precisely

- *r*_*i*_ is the intrinsic growth rate of primary producers (in [*time*]^-1^; *r*_*i*_ is positive for primary producers and null for other species);
- *G*_*i*_ is the growth term described in equation (2) below;
- *∈o_j_* is a conversion efficiency (dimensionless) which determines how much biomass eaten of resource *j* is converted into biomass of consumer *i*;
- *F*_*ij*_ is the functional response, i.e. the rate at which consumer *i* feeds on resource *j* (see equation (3) below; in [*time*]^-1^);
- *x*_*i*_ is the metabolic demand of consumer species *i* (in [*time*]^-1^); note that for basal species, metabolic demand is already taken into account in the intrinsic growth rate *r*_*i*_;
- *d*_*i*_ the natural mortality rate (in [*time*]^-1^).

#### Plant growth

We assumed a logistic growth for basal species:

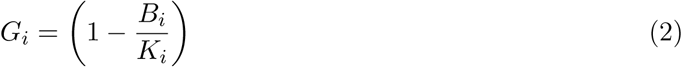

with *K*_*i*_ the carrying capacity of the environment for species *i* (in 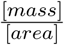)

##### Functional response

We used a multi-prey Holling-type functional response. The feeding rate of species *i* on species *j* is expressed as:

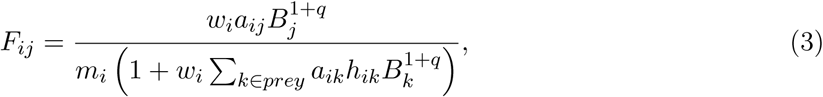

where:

- *w*_*i*_ is the relative consumption rate of predator *i* on its prey, which accounts for the fact that a consumer has to split its consumption between its different resources (dimensionless);
- *a*_*ij*_ is the capture coefficient in 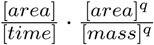(the attack rate here is 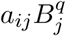which has the unit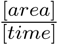
- 1 + *q* is the Hill-exponent, where the Hill-coefficient *q* makes the functional response vary gradually from a type II (q=0) to a type III (q=1) [52] (dimensionless);
- *h*_*ij*_ is the handling time in 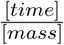.

#### 4.1.2 Introducing non-trophic interactions

We introduced in this model a number of non-trophic effects found to be frequent ones mentioned in the literature [3, 10, 21]. We made the relevant parameters of the trophic model become a function of the density of the species source of the effect [3, 18]. As a first approximation, we assume all such dependencies to have a similar, linear shape.

##### Competition for space

We can add for all species in the web a space-dependent term, *g*_*i*_, which affects their net growth rate:

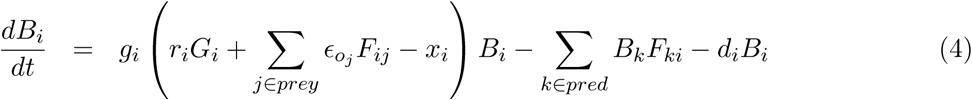

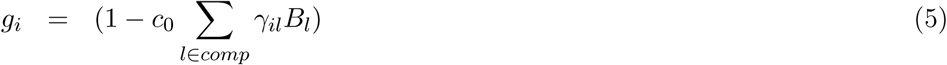

where *l* refers to all the species that potentially compete for space with each other (excluding intra-specific competition, i.e. *l* different from *i*), *c*_0_ is the overall intensity of competition for space and *γ_il_* is the strength of competition exerted by species *l* on species *i*, which is assumed to increase with the amount of space occupied by each individual of species *l* (see upcoming subsection on ‘Parameter values used’ in part 4.2). Note that the element *γ_il_* is zero if either *i* or *l* is non-sessile and even if both species are sessile it is non-zero only with a certain probability (see subsection 4.2, simulated networks). This makes competition for space asymmetric, as some species can have a large negative effect on others but are not negatively affected themselves. Also note that if *γ_il_* is null for all *l*, this equation (4) is identical to equation (1).

Competition for space was assumed to only operate if the net growth rate of the target species (i.e. first term in between brackets in equation (4) is positive (i.e.(*r*_*i*_*G*_*i*_ +Ʃ *_j∈ prey_* ∈*_oj_ F_ij_ - x_i_*)> 0). 0therwise *g*_*i*_ is set to 1.

##### Predator interference

We introduced predator interference in the feeding rate as follows:

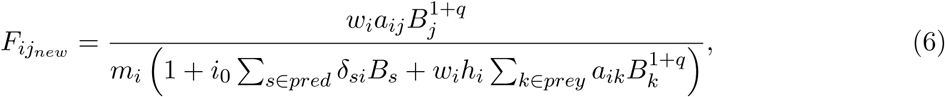

where *s* denotes all the predators of prey *j* and *δ_si_* is the strength of interference competition between predators *s* and *i* (Beddington-DeAngelis type, [12, 53]). Again, even if two predators share a prey, this term is non-zero only with a certain probability, and its values is assumed to depend on the differences of the body mass of the two predators (see upcoming subsection on ‘Parameter values used’ in part 4.2). The constant *i*_0_ is the overall intensity of predator interference.

##### Effects on mortality

A number of negative interactions lead to a decrease in the survival of the target species *i* (e.g. whiplash). Some species might also increase the survival of target species (e.g. improvement of local environmental conditions). We summarized these two types of effects on the mortality rate, *d*_*i*_ as follows:

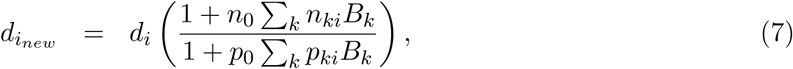

with *n*_0_ and *p*_0_ the overall intensity of negative and facilitative effects on mortality (i.e. resp. increase and decrease in mortality), and *n* and *p* the interaction matrices containing zeros and ones.

##### Refuge provisioning from predators

Refuge provisioning can happen in different ways: a species can protect another from abiotic stress (e.g. decreasing its mortality - see previous example) but a species can also protect another from its predator (e.g. affecting the attack rate of the predator). A refuge provision from species *k* to species *j* from its predators can be modeled as follows. For all the predator species *i* of *j*:

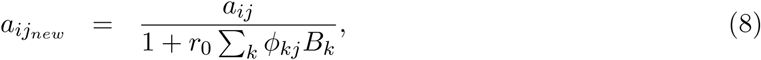

where *a*_*ijnew*_ tends to 0 in the presence of facilitators, *r*_0_ is the intensity of the refuge effect and *φ* is the interaction matrix containing zeros and ones.

### Effects on recruitment

Species may increase (e.g. habitat amelioration) the recruitment of new plants in the community. We therefore created the term *e*_*i*_ which is multiplied by the growth rate of the species:

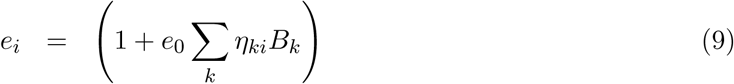

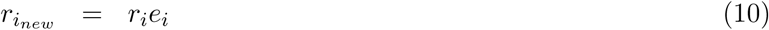

with *e*_0_ the overall intensity of negative and facilitative effects on recruitment, and *η* the interaction matrices containing zeros and ones. Note that this term only applied on plants.

### 4.2 Numerical simulations

#### Simulated networks

Trophic networks were generated with the niche model [51] starting with 100 species including a fixed number of 20 plants (primary producers) and a connectance of 0.06 (i.e. about 600 trophic links per network) [54]. Cannibalism was allowed but trophic networks containing cycles were discarded.

We imposed a fixed number of 33 sessile species in each network: for each species, we uniformly drew a ‘mobility’ trait with probability 0.2 for plants and 0.8 for other species, and we repeated the procedure until the network contained 33 sessile species.

The location of the non-trophic links was chosen randomly in the trophic web, but following a number of basic rules inspired from the Chilean data set [10, 21]. Competition for space was drawn between two sessile species, interference between two mobile predators that have at least one prey in common, refuge provisioning from a sessile species to a prey (i.e. a species that has at least one consumer), and recruitment facilitation from any species to a plant. Effects on mortality were drawn between any pair of species (i.e. no constraint). Non-trophic links were drawn only between different species because we focused on the role of inter-specific interactions. Simulations showing the effects of intra-specific competition for space and predator interference, both common in nature and well-studied in the literature, are shown in the Supplementary Materials (see supplementary Fig. A.5).

With the previous rules satisfied, the interaction probability (i.e. the probability for a non-zero element of the corresponding non-trophic interaction matrix) between two species was respectively set to 0.098, 0.15, 0.01, 0.01, 0.033 and 0.063 for competition, interference, increase and decrease in mortality, refuge provisioning and recruitment facilitation. These settings allowed to get, on average, 100 non-trophic links for any of the six types of non-trophic interactions (because of the imposed rules, the probabilities need to be different for each interaction type). This means that a simulation started with about 600 trophic and 100 non-trophic links of a given non-trophic interaction type. In the case of intra-specific interactions, the interaction probability was set to 1.

#### Simulations setup

In the first part (Fig. 1; see also supplementary Fig. A.1 and A.2), simulations were first run with trophic links only, and 100 trophic networks were selected in which no disconnected plant (i.e. with no consumer) was present at the end of the dynamics (as in [38]). For each of these trophic networks, we drew 100 non-trophic networks of each non-trophic interaction type (to study the effect of each type of non-trophic interaction individually). Again, we only kept the networks where no disconnected plant was present at the end of the dynamics with the non-trophic links. For each non-trophic interaction type, we started from a low non-trophic intensity value (0.5 and 10 for the negative and positive effects on mortality respectively, 0.1 for the others) and we linearly increased this value until a significant effect on diversity was observed compared to the case where there were only trophic interactions: 10% variation in species diversity, except for positive effects on mortality where reaching such an effect was not possible even for very high values of *p*_0_ (see dashed line in Fig. 1). With this procedure, we put all non-trophic interactions on equal footing, which allowed comparing their effect on outcome variables. This procedure allowed us to compute the slope of the effect of the non-trophic interaction intensity on final species diversity using a linear regression, which is an indicator of the strength of the non-trophic interaction. The slope of this regression is displayed on top of each panel of Fig. 1.

In the second part (Fig. 2, 3; see also supplementary Fig. A.4), decrease in mortality were discarded because of their lack of significant effect on species diversity. For the other five non-trophic interaction types, we selected 1000 simulated trophic networks (again with no disconnected plants), and for each of these networks, we repeated the following procedure 100 times: we drew five non-trophic networks (one per non-trophic interaction type), with one quarter of the previous interaction probabilities for the four negative non-trophic interaction types. We uniformly drew the non-trophic interaction intensities in the same range of values as in the first part (see above). Hence, in this set of simulations, we started with about 600 trophic links, 100 positive (facilitation for recruitment) and 100 negative non-trophic links (about 25 for each of the four types: competition for space, interference, refuge and increase in mortality).

In the same vein, we also did another set of simulations in which the number of links with fixed non-trophic intensities varied (those leading to 10% variation in species diversity; *i*_0_ = 2.75, *r*_0_ = 0.9, *c*_0_ = 0.3 and *e*_0_ = 0.5 *n*_0_ = 6.5) (Fig. A.3). We simulated 100 trophic networks with no disconnected plants and, for each trophic network, we drew 100 non-trophic interaction networks of each non-trophic interaction type of varying size: by modulating the previously mentioned interaction probabilities, we created setups with 100 positive links and 100 negative links (100 of one type or about 25 for each of the four types), 50 and 100, or 100 and 50 respectively. Again we only retained the network with no disconnected plant at the end of the dynamical simulations with trophic and non-trophic links.

#### Simulation runs

We used the GNU Scientific Library (https://www.gnu.org/software/gsl/) solver with the embedded Runge-Kutta-Fehlberg (4,5) method. For each network, numerical simulations were run until steady state was reached (we set a maximum time *t*=1000 which we observed to be sufficient). During the dynamics, we set a species to extinction when its biomass was very small (<1e-6). To fairly compare results with and without non-trophic interactions, we used the same initial conditions in both cases. At steady state, we measured: diversity, i.e. the number of surviving species (biomass>=1e-6), total biomass (sum of the biomass of all surviving species at steady state) and total production (sum of all the positive terms of the dynamical equations of all surviving species). We also evaluated the normalized ratio of each of these metrics in the case with and without non-trophic interactions. For instance, we call *diversity ratio* (in particular in the legends of the figures) the difference between the diversity with and without non-trophic interactions divided by the diversity without non-trophic interactions.

#### Software availability

All simulations were performed with our C++ program dynaweb which is licensed under the GNU Public Licence and freely available upon request.

### Parameter values used

- ∈_0*j*_ was set to 0.45 if the resource *j* is a plant and to 0.85 otherwise;
- *r*_*i*_ is the intrinsic growth rate of plant species with 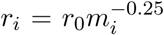if species *i* is a plant, and *r*_*i*_ = 0 for consumer species [55]; The unit of *r*_*i*_ is [*time*]*^-1^* and *r*_0_ is a scaling parameter which is the same for all species and has the unit 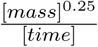(it defines the time scale of the system). *r*_0_ = 1.
- *x*_*i*_ is the metabolic demand of species *i*. If *i* is not a plant 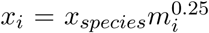 with *x*_*species*_ = 0.314
- *d*_*i*_ the natural mortality of species *i* is assumed to be 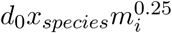 with *d*_0_ = 0.1 and *x*_*species*_ = 0.138 if *i* is a plant and 0.314 otherwise.
- *K*_*i*_ the carrying capacity of the environment for species *i (in 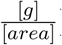)*; 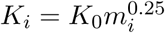 with *K*_0_=1 (in case of logistic growth).
- *w*_*i*_, the relative consumption rate; it is defines as 1*/*(number of resources of species *i*);
- *a*_*ij*_ is the capture coefficient, such that if *i* and *j* are both mobile species: 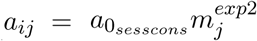 [56] with *exp*1 = 0.45, *exp*2 = 0.15 and *a*_0_ = 10; if *i* is sessile and *j* is mobile: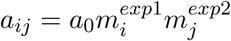 with 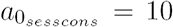; if *i* is mobile and *j* is sessile: with 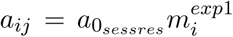with 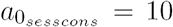
- 1 + *q* is the Hill-exponent; 1 + *q* = 1.2.
- *h*_*ij*_ is the handling time in [*time*], with 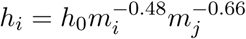and *h*_0_ = 0.4 [49, 57].
- *γ_ij_*, the effect strength of competition for space of species *j* on species *i*, is assumed to depend on body mass such that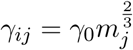, with *γ*_0_ = 1.
- *δ_ij_* is the term of interference between predator *i* and predator *j*; we assume that the more similar the body masses of the two predators, the stronger the interference between them, such that 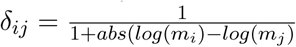
- We make the non-trophic interaction intensities vary in the following ranges : 0.1 ≤ *c*_0_ ≤ 0.4, 0.25 ≤ *i*_0_ ≤ 4, 1 ≤ *n*_0_ ≤ 7, 5 ≤ *p*_0_ ≤ 20, 0.1 ≤ *r*_0_ ≤ 1.2, 0.1 ≤ *e*_0_ ≤ 1.2
- Body mass of each of the species in the networks were determined based on the trophic level: 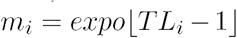, with *expo* = 2.6, *TL*_*i*_ the trophic level of species *i* and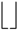 indicating that the term in the middle in rounded down to the nearest integer.

## ASupplementary figures

**Figure A1:**
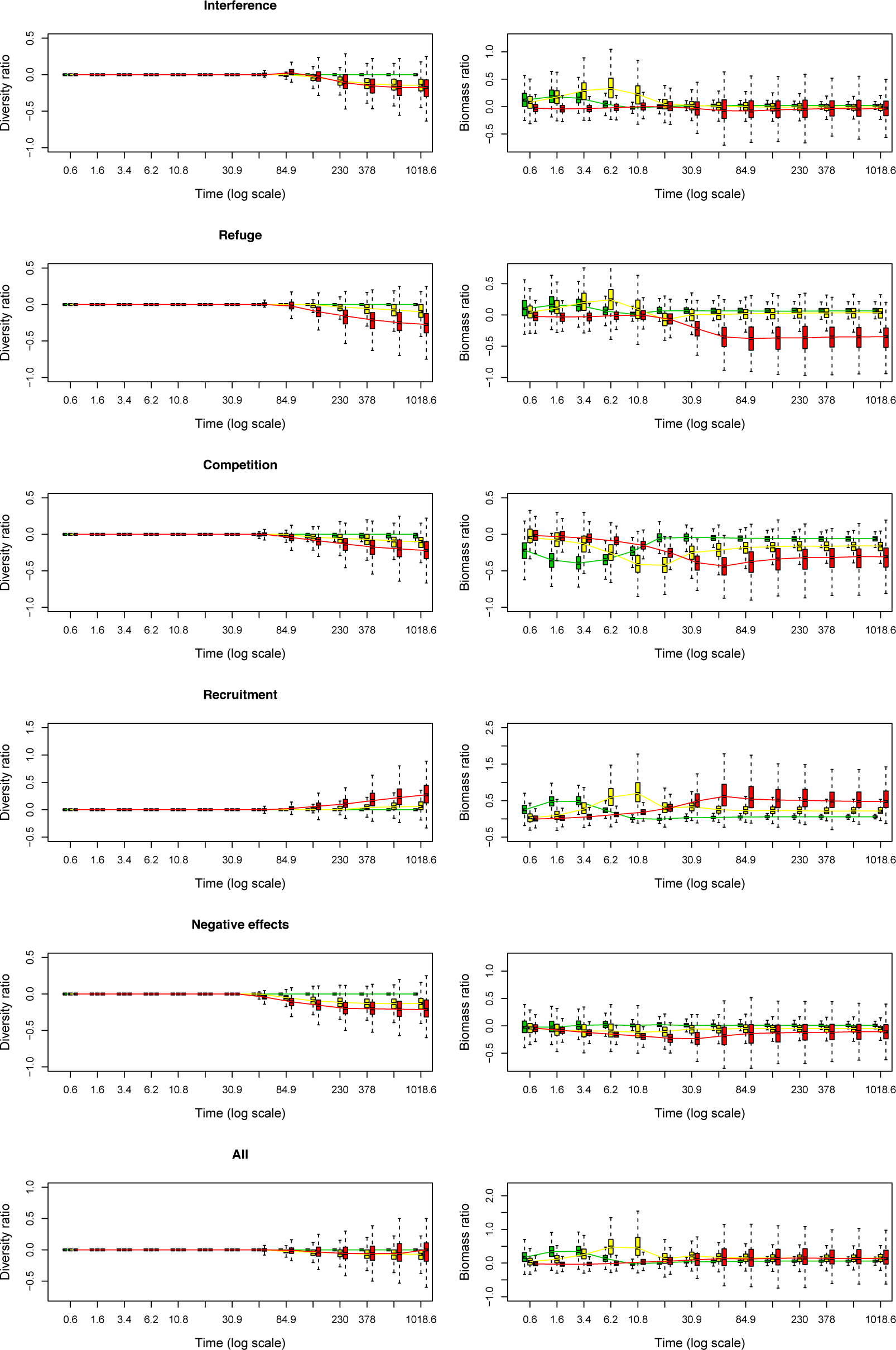
Supplementary Figure. For the main NTI types (5 first rows) and the NTI all together (6th row), relative change in species diversity (left column) and biomass (right column) per trophic level through time during 500 simulations. Each simulation starts with 600 trophic links and 100 non-trophic links. Note that on the *x*-axis, time is on a log-scale. Green: TL=1, yellow: TL between 2 and 3, red: TL> 3.

**Figure A2:**
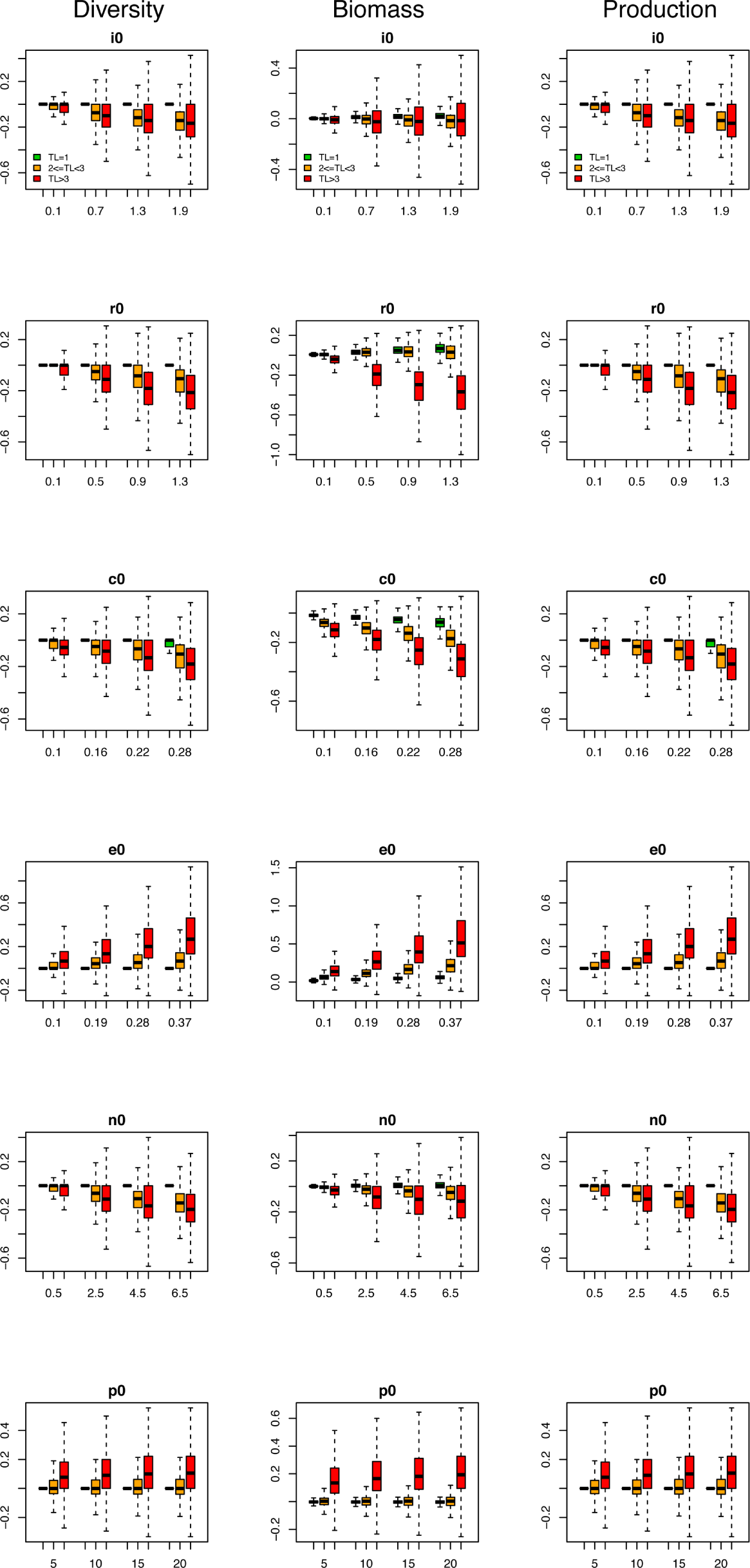
Supplementary Figure. For each NTI type (6 rows), similarly to Fig. 1, each of the measured variables (3 columns) as a function of the intensity of the NTI, by trophic level. Note that each NTI has its own y-axis. Green: TL=1, yellow: TL between 2 and 3, red: TL> 3.

**Figure A3:**
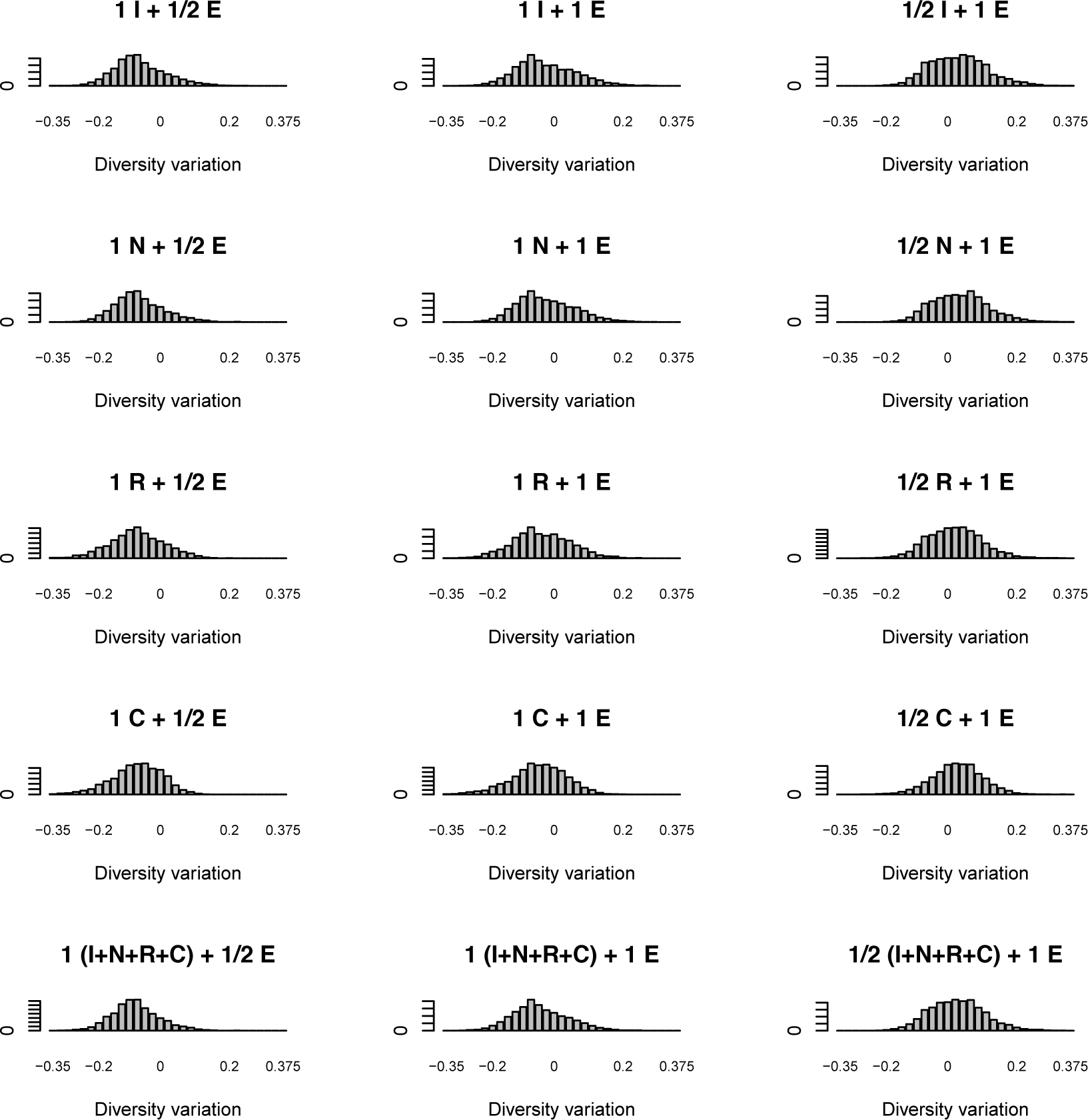
Supplementary Figure. Frequency of simulations leading to a given variation in species diversity (in the case with compared to without NTI links). The intensity of each NTI type is now fixed and we vary the relative number of links of different NTI types when put together. The left column correspond to situations where there are twice more negative than positive links, the middle column shows results with equal number of positive and negative links, and the right column correspond to cases where there are twice more beneficial than detrimental links. The panels correspond to different combinations of NTIs: I for interference, E for recruitment facilitation, N for negative effects on mortality, R for refuge, C for competition. At fixed intensity and with equal number of links, negative links tend to take over (slightly). There are configurations of relative abundance of the different types of NTIs in which positive and negative effects can balance each other.

**Figure A4:**
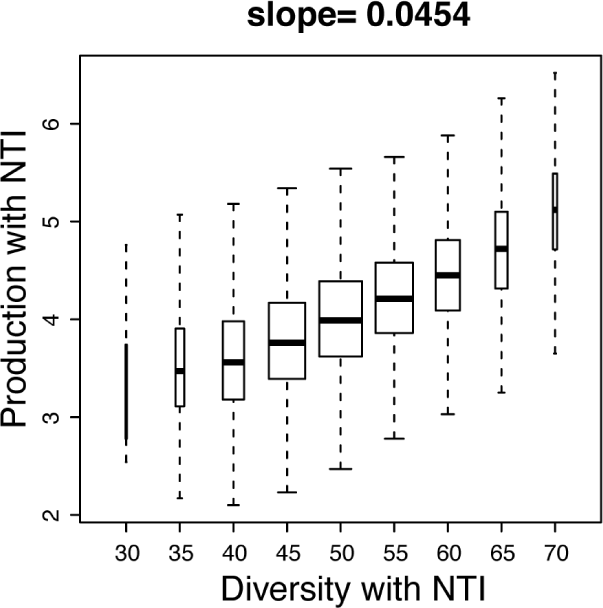
Supplementary Figure. Same as Fig. 3 B but for production instead of biomass. Variation in production as a function of species diversity in networks with NTIs. The slope obtained by linear regression of the relationship is indicated on top of the panel.

**Figure A5:**
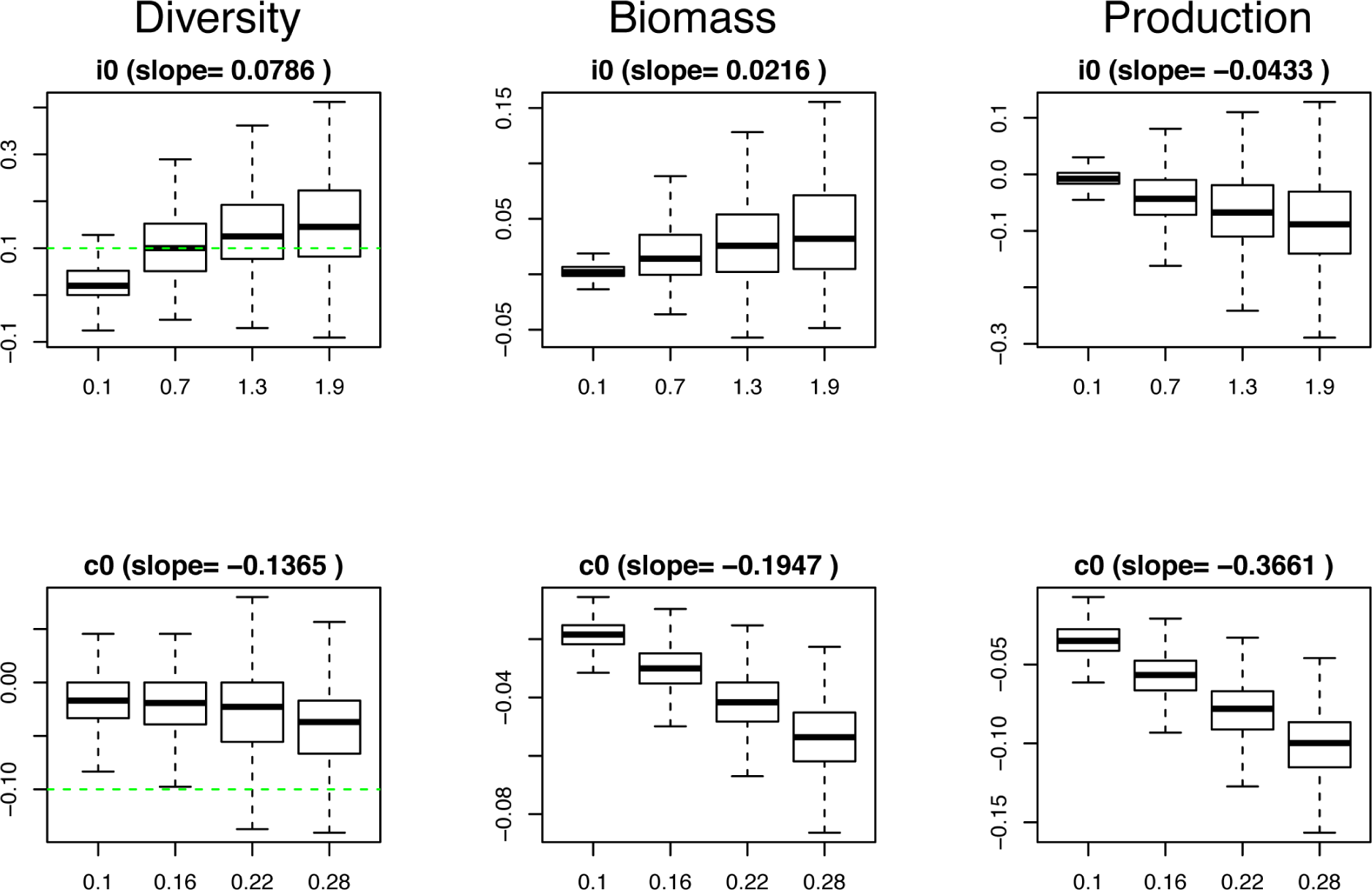
Supplementary Figure. Same as Fig. 1 but for intra-specific interference (top row) and intra-specific competition for space (bottom row).

